# Selective translation by alternative bacterial ribosomes

**DOI:** 10.1101/605931

**Authors:** Yu-Xiang Chen, Zhi-yu Xu, Xueliang Ge, Suparna Sanyal, Zhi John Lu, Babak Javid

**Affiliations:** Centre for Global Health and Infectious Diseases, Collaborative Innovation Centre for the Diagnosis and Treatment of Infectious Diseases, Tsinghua University School of Medicine, Beijing, China; MOE Key Laboratory of Bioinformatics, Center for Synthetic and Systems Biology, School of Life Sciences, Tsinghua University, Beijing, China; Department of Cell and Molecular Biology, Uppsala University, Sweden

**Author notes:** These authors contributed equally to this work (Yu-Xiang Chen, Zhi-yu Xu). to whom correspondence should be addressed: ZJL, BJ (lead contact).

**Keywords:** alternative ribosomes, specialized ribosomes, mycobacteria, ribosome profiling

## Abstract

Alternative ribosome subunit proteins are prevalent in the genomes of diverse bacterial species but their functional significance is controversial. Attempts to study microbial ribosomal heterogeneity have mostly relied on comparing wild-type strains with mutants in which subunits have been deleted, but this approach does not allow direct comparison of alternate ribosome isoforms isolated from identical cellular contexts. Here, by simultaneously purifying canonical and alternative RpsR ribosomes from *Mycobacterium smegmatis*, we show that alternative ribosomes have distinct translational features compared with their canonical counterparts. Both alternative and canonical ribosomes actively take part in gene translation, although they translate a subset of genes with differential efficiency as measured by ribosome profiling. We also show that alternative ribosomes have a relative defect in initiation complex formation. Our work convincingly confirms the distinct and non-redundant contribution of alternative bacterial ribosomes for adaptation to hostile environments.

**Significance Statement:** Many organisms, including most bacteria code for multiple paralogues of some ribosomal protein subunits. The relative contribution of these alternative subunits to ribosome function and gene translation is unknown and controversial. Furthermore, many studies on alternative ribosomes have been confounded by isolation of alternative and canonical ribosomes from different strains and/ or different growth conditions, potentially confounding results. Here, we show unequivocally that one form of alternative ribosome from *Mycobacterium smegmatis* actively engages in gene translation, but its translational profile from an identical cellular context is subtly different from canonical ribosomes. Given the prevalence of alternative ribosomal genes in diverse organisms, our study suggests that alternative ribosomes may represent a further layer of regulation of gene translation.

## Introduction

Ribosomes are the macromolecular machines that translate the genetic code into functional proteins (1–3). Shortly after their discovery and role in gene translation, the “one gene, one mRNA, one ribosome” hypothesis was proposed (4), but was quickly disproven (5). This led in turn to the “homogeneity hypothesis” for ribosomes: that all ribosomes were identical macromolecules, that would translate all mRNAs with equal efficiency (6). However, observations that ribosome composition varied according to environmental conditions and other variables, immediately challenged the homogeneity hypothesis (7). But until recently, the functional significance of ribosome heterogeneity has not been well understood.

In eukaryotic systems, there is increasing evidence for the role and importance of ribosomal heterogeneity in fundamental physiology, development and disease, in systems from budding yeast, to organelles and animals (8–11). In particular, specialized ribosomes have been shown to be important for vitamin B12 transport and cell-cycle components (12), hematopoiesis (13) and development (14). Mutations in ribosomal proteins, or haploinsufficiency of their coded genes have been implicated in a number of disorders such as the ribosomopathies (9, 15). Our understanding of ribosomal heterogeneity in bacteria is less advanced (16). Using high-fidelity mass spectrometry of intact 70S ribosomes from *Escherichia coli* revealed considerable heterogeneity, such as the presence or absence of the stationary-phase-induced ribosomal-associated protein, SRA (17). Other studies implicating specialized bacterial ribosomes (18, 19) have recently been questioned (20–23). Nonetheless, it is intriguing to speculate that specialized bacterial ribosomes, generated in response to environmental stressors, would result in altered translation, which in turn might allow adaptation to the stressful environment (16).

Specialized ribosomes could be generated via changes in the stoichiometry of canonical ribosomal components, association of accessory ribosomal proteins, modification of ribosomal rRNA or incorporation of alternative ribosomal subunits, coded by paralogous or homologous genes (reviewed in (9, 24)). Most eukaryotic organisms code for paralogues of ribosomal subunit genes, and around half of sequenced bacterial genomes from one study included at least one ribosomal protein paralogue (25). One well-characterized group of paralogous ribosomal proteins are those that lack cysteine-rich motifs (C-) compared with canonical cysteine-containing homologues (C+). C- paralogues have been described in many bacteria, including mycobacteria (26–28). In mycobacteria, the conserved alternative C- ribosomal subunits are coded in a single operon regulated by a zinc uptake regulator (zur), which represses expression in the presence of zinc ions (29), and has led to the suggestion that the function for C-/C+ ribosomal subunit paralogues is to allow for dynamic storage of zinc (30–32). Furthermore, although these alternative mycobacterial ribosomal proteins (AltRPs) have recently been demonstrated to incorporate into assembled ribosomes, it has been suggested that they form non-translating hibernating ribosome complexes due to exclusive association of a hibernation factor with alternative ribosomes (33). Here, we demonstrate convincingly using a fully reconstituted mycobacterial translation system that ribosomes containing the alternative subunit RpsR2 (AltRpsR) actively translate. Furthermore, the translational landscape as measured by ribosome profiling of these alternative mycobacterial ribosomes are distinct from those generated by canonical ribosomes purified from the same cell and are characterized by a 5’ polarity shift suggesting that alternative bacterial ribosomes may provide a further layer of regulation during nutrient stress.

## Results

### Alternative mycobacterial ribosomes engage in gene translation

Mycobacteria encode for several alternative C- ribosomal proteins, four of which are conserved throughout the genus and are encoded within one operon (27, 33) (Fig. 1A). We wished to biochemically characterize alternative ribosome (AltRibo) function by purifying ribosomes containing alternative or canonical subunit proteins. We chose to C-terminally tag RpsR2 (AltRpsR) with 3xFLAG, since of the four alternative ribosome genes within the operon, the C-terminus of RpsR showed the greatest sequence variation (Fig. 1A), and we reasoned that it therefore was most likely to tolerate tagging. Immunoprecipitation of C-terminally-FLAGx3-tagged alternative RpsR2 co-precipitated all ribosome subunits (Table S1 and Fig. S1A, B), suggesting that purification of alternative and canonical ribosomes could be achieved by tagging the two isoforms of RpsR.

**Figure 1.**
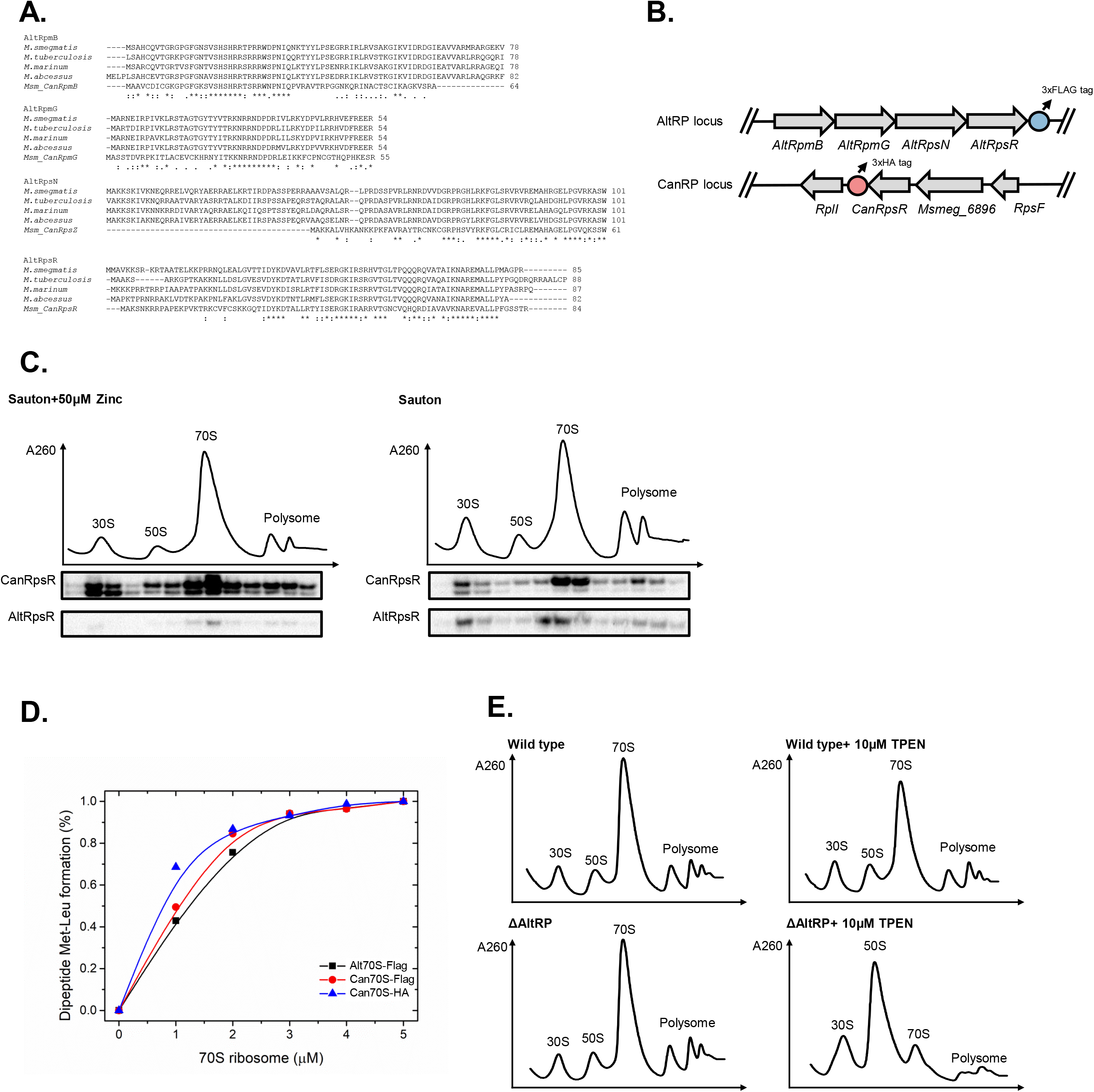
Alternative mycobacterial ribosomes engage in gene translation. (A) Sequence alignments of the alternate ribosome proteins from 4 mycobacterial species. The corresponding canonical RP from *M. smegmatis* is shown below the alternate genes for comparison. (B) Cartoon representing the C-terminus affinity tagging of the native CanRpsR and AltRpsR native loci. (C) Subunit profiling of the strain depicted in (B) cultured in either Sauton medium or Sauton medium supplied with 50μM. The corresponding 30S, 50S, 70S and polysome fractions are indicated. Each fraction was blotted with anti-FLAG (AltRpsR) and anti-HA (CanRpsR) antibody. (D) Di-peptide formation of AltRibos vs CanRibos with a fully reconstituted mycobacterial translation system (34). Peptides were detected and analyzed by high performance liquid chromatography. (E) Subunit profiling of wild type *M. smegmatis* and AltRP knockout strain following plating on 7H10 agar +/- supplementation with10 μM TPEN. The corresponding 30S, 50S, 70S and polysome fractions are indicated.

To investigate alternative (AltRibo) and canonical (CanRibo) RpsR ribosome function within the same cellular context, we constructed a strain of *M. smegmatis* in which both RpsR subunits had affinity tags inserted at their C-terminus, at their native loci on the mycobacterial chromosome (Fig. 1B and see Methods). In keeping with prior studies (28, 34), growth of mycobacteria in relatively zinc-deplete Sauton’s medium upregulated AltRpSR expression and supplementation with zinc restored CanRpsR expression (Fig. S2). We grew the dual-tagged *M. smegmatis* strain in both zinc-replete and relatively zinc-deplete conditions, purified ribosomes and subjected them to subunit profiling. Under both conditions, translating ribosomes (polysome fraction) could clearly be identified (Fig. 1C). Immunoblotting against AltRpsR-FLAG and CanRpsR-HA confirmed that zinc depletion upregulated AltRpsR expression. Importantly, a significant proportion of polysomes isolated from growth in zinc-depleted medium were AltRibos (Fig. 1C), suggesting that AltRibos are resident on translating polysomes and supporting their active role in gene translation. We further subjected affinity purified AltRibos and CanRibos to biochemical analysis in a mycobacterial cell-free translation system (34). AltRibos were able to synthesise di-peptides at similar efficiency to CanRibos (Fig. 1D), unequivocally confirming that AltRibos are capable of gene translation.

We also constructed a strain in which the alternative subunit operon was knocked out, ΔARP. This strain grew poorly with zinc depletion (Fig. S3), confirming prior results. We wondered whether this might be because under conditions of zinc depletion, CanRibos are less proficient at gene translation? We performed subunit profiling on both wild-type and ΔARP *M. smegmatis*, scraped from agar plates grown both on standard medium (7H10-agar) and plates supplemented with the specific zinc chelator N,N,N,N’-tetrakis(2-pyridinylmethyl)-1,2-ethanediamine (TPEN) at a concentration of TPEN that did not interfere with bacterial growth (Fig. S4). Although the polysome fraction was preserved in both strains when isolated from standard growth conditions, the polysome fraction of ΔARP was significantly smaller under zinc depletion (Fig. 1E), suggesting that canonical ribosomes may translate less effectively under zinc depletion, and that alternative-RpsR ribosomes are likely to be the major translating fraction under those conditions.

### Alternative mycobacterial ribosomes have distinct translational profiles compared with canonical ribosomes

The dual-RpsR-tagged strain (Strain A) permitted us to interrogate selective ribosome translation profiles from the same cellular population, which would eliminate potential differences due to mRNA abundance or other confounding factors (35, 36). We grew the strains in relatively zinc-replete conditions, which allowed cultures to grow to greater density for ribosome isolation prior to ribosome profiling (35, 37–39). Under these conditions, the majority of ribosomes were CanRibos. To exclude the formal possibility that observed differences were due to the use of the affinity tags, we generated a second dual-tagged strain in which the affinity tags were swapped (Strain B – Fig. 2A). Both strains were similarly regulated by zinc (27, 33) (Fig. S2), suggesting they had comparable physiology.

**Figure 2.**
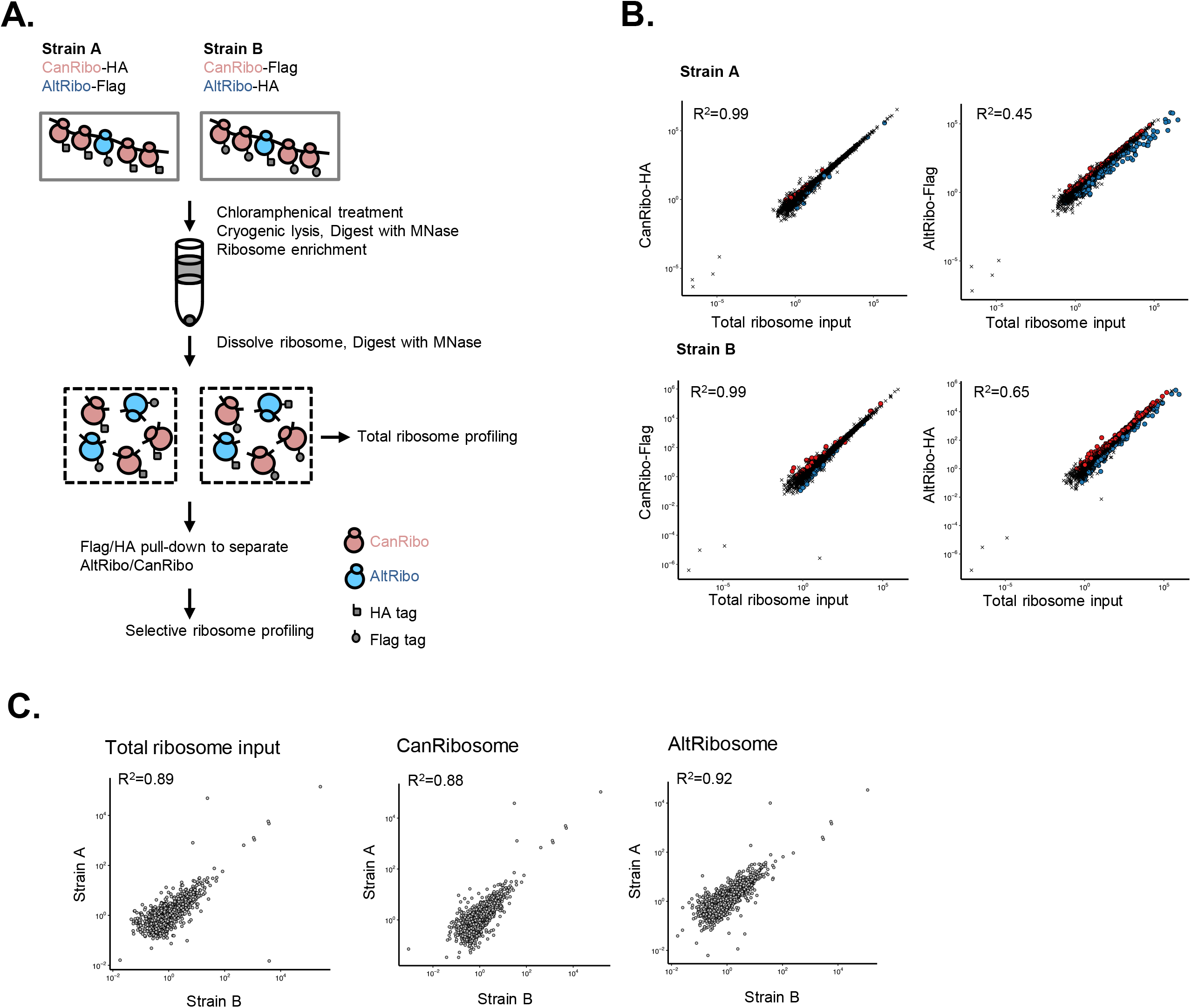
Alternative ribosomes have distinct translation profiles. (A) Cartoon outlining work-flow for performing selective ribosome profiling in the two RpsR-tagged strains – Strain A and Strain B (and see Methods). (B) Scatter plot showing selective ribosome profiles compared with total ribosome input in Strain A and Strain B. Axes represent reads per kilobase per million mapped reads (RPKM). Blue and red dots represent down- and up-regulated reads respectively. R^2^ represents Pearson correlation coefficient. (C) Correlation of translation efficiency of total ribosome, selective CanRibo and selective AltRibo inputs in Strain A compared with Strain B. R^2^ represents Pearson correlation coefficient.

Position analysis of aligned reads revealed a clear 3 nucleotide periodicity (Fig. S5), confirming that reads were derived from translating ribosomes for both AltRibos and CanRibos, and showed a high degree of reproducibility between replicates (Fig. S6A). Comparison of translational profiles confirmed that CanRibo-derived reads were highly correlated with total ribosome input, consistent with the fact that the majority of ribosomes under the tested condition were CanRibos. However, reads generated from AltRibos were less well correlated, suggesting that translational profiles from AltRibos were distinct from CanRibos (Fig. 2B). Regardless of affinity tag, CanRibo- and AltRibo-derived reads were highly correlated with each other when comparing translational efficiency profiles generated by Strain A and Strain B (Fig. 2C), following correction for mRNA expression (Fig. S6B). These data together strongly support that AltRibos and CanRibos generate distinct translational landscapes.

Our data suggested that alternative ribosomes had intrinsic differences in gene translation compared with canonical ribosomes. We performed differential expression analysis on AltRibo- and CanRibo-generated ribosome profile datasets, and compared these with the total ribosome input. Pull-down efficiency of HA-tagged subunits was less efficient than FLAG-tagged subunits. Although this was not an issue in Strain A, where the highly abundant CanRibos were HA-tagged, it meant that the number of reads from the HA-tagged AltRibos from Strain B were lower, and as such, fewer genes were differentially translated (up- or down-) in AltRibos isolated from Strain B. As expected, there were few differences between CanRibos and total ribosomes. By contrast, there were a large number of genes that were translated with either decreased or increased preference by AltRibos compared with total ribosome input (Fig. S7A, B). Of note, there was significant enrichment in under-represented translated genes in AltRibos isolated from Strain A and B. (Fig. S7B, C).

### Alternative ribosomes have polarized read distribution a relative initiation defect

Our analysis of differences in translational profiles between canonical and alternative mycobacterial ribosomes revealed that for translation of certain genes, there were clear differences in the distribution of sequencing reads along the gene length between the two samples, for example for translation of *Msmeg_0363* (Fig. 3A). To characterize these differences on a genome-wide basis, we calculated the polarity score to determine the distribution of aligned reads for each coding gene (40). Polarity scores between −1 and 0 represent read accumulation at the 5’ end of the coding region, and between 0 and +1 represent read accumulation at the 3’ end of the coding region (see Methods). Comparing the polarity score of AltRibos with CanRibos and total ribosome input from both Strain A and Strain B, there was a clear polarity shift towards a 5’ read accumulation in alternative ribosomes (Fig. 3B).

**Figure 3.**
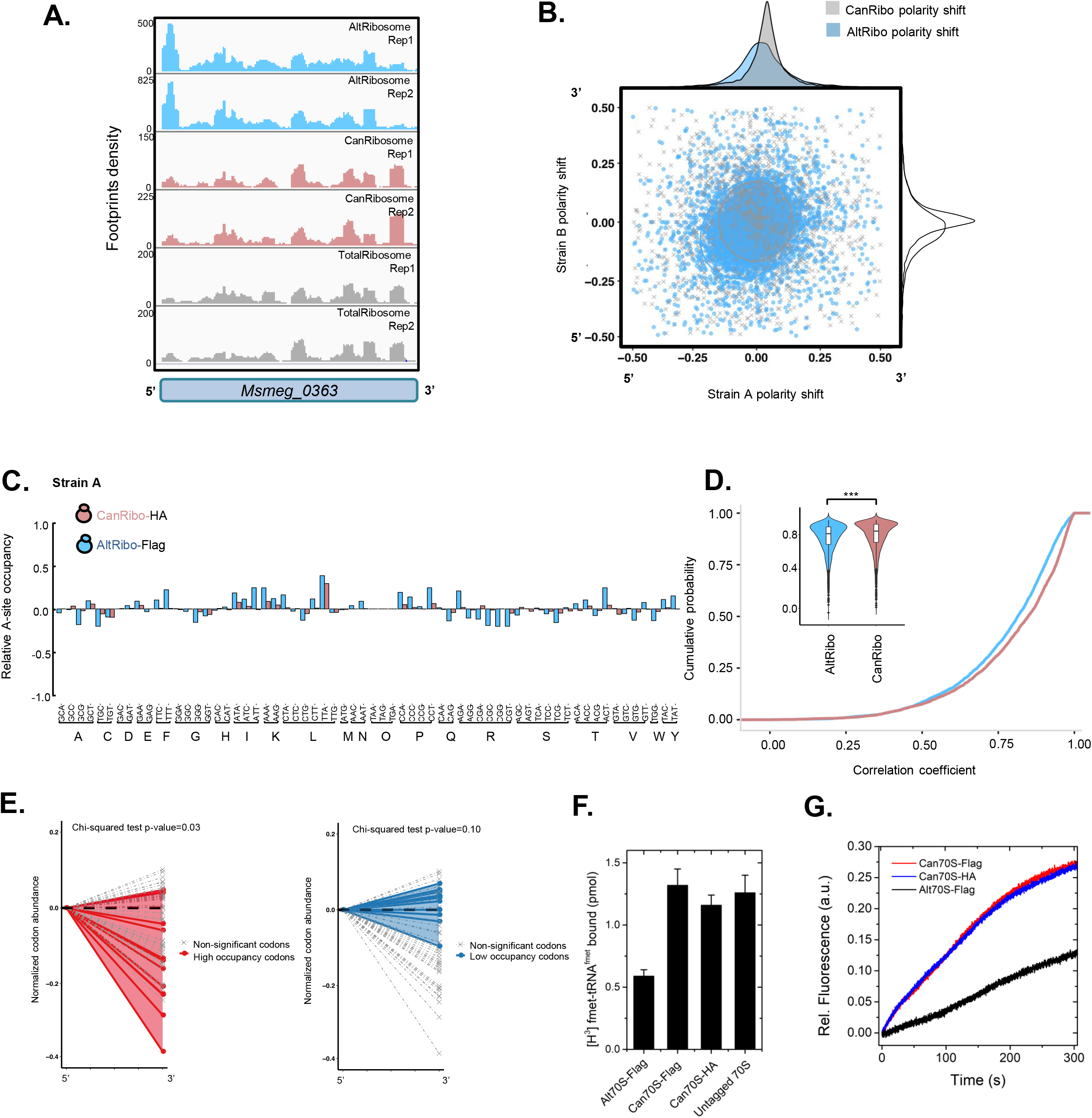
Alternative ribosomes have 5’ read accumulation and a relative initiation defect compared with canonical ribosomes. (A) Normalized reads of each ribosome input for the gene *Msmeg_0363* were plotted along gene length using IGV software (see Methods). Reads from AltRibos show a clear 5’ accumulation compared with reads generated from CanRibos or total ribosomal input. (B) Shifts in polarity scores for the entire genome generated from selective AltRibo and CanRibo reads from Strains A and B – blue dots represent shifts calculated from AltRibos and gray squares represent shifts calculated from CanRibos. The overall polarity score shifts are illustrated as smoothed curves above and to the right of the plot. A negative score indicates 5’ read accumulation and a positive score indicates a 3’ read accumulation. (C) The relative A-site codon occupancy for all 61 sense codons for AltRibos and CanRibos in Strain A compared with total ribosome input. (D) Cumulative probability and violin (inset) plots for correlation between codon occupancy for AltRibos and CanRibos compared with total ribosome input from Strain A. *** p<0.001 by Student’s t-test. (E) The relative codon frequencies of the 5’ 50 codons and 3’ 50 codons for each ORF were analyzed. Comparing AltRibo high occupancy codons (left panel) and low occupancy codons (right panel) with non-significantly enriched codons, there was significant enrichment of high occupancy codons at the beginning of ORFs, and a trend towards enrichment of low occupancy codons at the end of ORFs by chi-squared test. (F) The extent of initiation complex formation by AltRibos compared with CanRibos measured by retention of ^3^H-fMet count on a nitrocellulose filter. (G) The kinetics and extent of binding of the initiator tRNA to the mRNA programmed 70S ribosome (AltRibo vs CanRibo) followed with BODIPY fluorescence.

Translation elongation does not proceed at a uniform pace (41–43). We investigated whether alternative and canonical ribosomes differed in codon-reading rates. AltRibos from either Strain A or Strain B showed similar, though not identical codon usage pattern across all 61 sense codons (Fig. 3C, Fig. S8A). Comparing CanRibos with AltRibos with total ribosome input across all coding genes, there were significant differences in codon usage between the two ribosome isoforms in both Strain A and Strain B (Fig. 3D and Fig. S8B). From our analysis of all 61 sense codons in both strains, 12 showed relatively high occupancy in AltRibos compared with total ribosomes, and 12 showed relatively low occupancy (Fig. S8C). We wished to determine whether altered codon usage might be associated with the observed polarity differences between AltRibos and CanRibos. We considered the first and last 50 codons of all coding genes and compared the relative abundance of the 12 high- and 12 low-occupancy codons in these regions. For the 12 high occupancy codons, 10 were relatively enriched at the 5’ end of coding regions, and for the 12 low occupancy codons, 8 were relatively enriched at the 3’ end of coding regions (Fig. 3E), potentially suggesting a mechanism for the observed polarity differences between alternative and canonical ribosomes. Furthermore, although purified AltRibos formed 70S initiation complexes, they appeared to do so at slightly lower efficiency compared with CanRibos (Fig. 3F), suggesting another potential mechanism for the observed 5’ polarity shift. The lower efficiency in 70S initiation complex formation could be due to impaired binding of the initiator tRNA as demonstrated by BODIPY-Met-tRNA^fMet^ binding to the ribosome monitored by increase in BODIPY fluorescence (Fig. 3G).

## Discussion

Together, our studies demonstrate that alternative mycobacterial ribosomes incorporating C- paralogues are not only competent for translation, but that these alternative ribosomes have distinct translational landscapes compared with their canonical counterparts. Although the ribosome is primarily an RNA machine (2), multiple studies have shown that alterations in ribosomal subunit proteins may have profound impacts on aspects of translation function (12, 36, 44, 45). We leveraged the power of ribosome profiling and affinity purification of specialized ribosomes to show that alternative RpsR mycobacterial ribosomes have altered codon usage compared with ribosomes containing the canonical RpsR paralogue, as well as a relative 5’ polarity shift in translation of open reading frames. It should be noted that due to our tagging and purification strategy, we were primarily comparing RpsR canonical and alternative ribosomes. Although due to their differential expression, we would expect that in high and low zinc environments, alternative and canonical ribosomes would contain mostly the entire canonical or alternative subunit proteins, in environments where expression of all subunits are sustained (such as our experimental conditions), multiple combinations of ‘alternative’ ribosomes could co-exist, and our data are unable to resolve between these. Ribosomal profiling of specialized ribosomes had previously demonstrated specific functions of heterogeneous ribosome populations in stem cells and for the translation of mitochondrial proteins in eukaryotic cells (12, 44). In bacteria, ribosome profiling of trigger factor associated with ribosomes demonstrated the requirement of trigger factor particularly for the translation of outer-membrane proteins in *E. coli* (36), but ribosome profiling had not been employed for ribosomes incorporating ribosomal protein paralogues.

There have been relatively few studies of bacterial ribosomal heterogeneity when compared with eukaryotic systems (16). However, there is evidence that ribosomal heterogeneity may be just as pervasive in bacteria as eukaryotes (46, 47). Mycobacteria, for example, have very distinct translational features compared with the model *E. coli* system (39, 48, 49). Recent structural information about the mycobacterial ribosome identified new ribosomal protein subunits, but these new subunits, bL37 and bS22 were not present in all ribosomes identified by cryo-electron microscopy (34, 50, 51).

Many diverse bacterial species code for C- ribosomal subunit paralogues (25), but their distinct translational functions, if any, have not been fully characterized. A recent examination of the mycobacterial C- paralogues that are the subject of this study suggested that the C- ribosomes are hibernating and do not actively engage in gene translation (33, 52). Those findings are in contradistinction with our own. By both biochemical analysis and ribosome profiling, our data support a role for translating AltRibos, even under conditions of zinc depletion. Ojha and colleagues suggested an exclusive association of a ribosomal hibernation promoting factor (*Msmeg_1878* and named MPY by them) with C- ribosomes under conditions of zinc depletion. It is possible that use of supra-physiological (1mM) zinc in culture medium led to lack of binding of MPY to canonical (C+) ribosomes under those conditions (33). However, our isolation of ribosomes from *M. smegmatis*-ΔAltRP (where C- ribosomes would be absent by definition) by standard methods in standard growth medium (Supplementary Table 1, Uniprot accession number A0QTK6) showed association of CanRibos with MPY and demonstration that canonical ribosomes are less competent at polysome formation under zinc depletion (Fig. 1E), as well as a previous study of *M. smegmatis* grown under conditions where C- ribosomes would not have been predominant (53) suggest that MPY is not exclusively associated with C- ribosomes.

Our data suggest that alternative ribosomes and canonical ribosomes have differences potentially in both translation initiation and codon usage – both of which may contribute to the relative 5’ polarity shift of reads in alternative ribosomes. Multiple mechanisms have been proposed for observed differences in elongation rates (54). Decoding of modified tRNA bases (42) and differential affinity of specific ribosome isoforms for aminoacyl-tRNAs (41) also contribute to altered rates of gene translation. Of note, RpS18 (of which AltRpsR and CanRpsR are the two mycobacterial isoforms) is part of an mRNA “nest”, which contributes to preinitiation binding of mRNA to the 30S subunit (55), and it is tempting therefore to speculate that differences in RpsR protein sequence may contribute the observed differences in initiation complex formation. Our study reveals not only that alternative mycobacterial ribosomes actively participate in gene translation, they but also provides evidence that gene translation by specialized bacterial ribosomes contribute to environmental adaptation.

## Materials and Methods

### Bacterial strains and culture

Wild-type *M. smegmatis* mc^2^-155 (56) and derived strains were grown in either Middlebrook 7H9 media supplemented with 0.2% glycerol, 0.05% Tween-80, 10% ADS (albumin-dextrose-salt) with appropriate antibiotics or Sauton’s medium (0.05% KH_2_PO_4_, 0.05% MgSO_4_·7H_2_O, 0.2% citric acid, 0.005% ferric ammonium citrate, 6% glycerol, 0.4% asparagine, 0.05% Tween, pH 7.4) (27). Zinc sulfate at indicated concentrations was added to Sauton medium as per specific protocols. If not otherwise noted, bacteria were grown and maintained at 37°C with shaking.

### *M. smegmatis* AltRP knock-out strain and complemented strain construction

In normal conditions (High Zinc), the AltRP operon is not required for growth. A double crossover strategy was used to construct an unmarked AltRP knockout strain as previously described (57). Sequences 1,000bp upstream and downstream of the operon were cloned and ligated into the P2NIL suicide vector. The lacZ and SacB gene were cloned from pGOAL17 and ligated to P2NIL containing upstream and downstream 1,000bp region to generate the AltRP deletion construct. 2μg plasmid were transformed to wild-type *M. smegmatis* competent cells by electrophoresis and selected on LB plates supplemented with 25μg/ml kanamycin and 50μg/ml X-gal. Blue colonies were screened for single crossover recombinants and further verified by PCR. Correctly identified recombinants were further plated on LB plates containing X-gal and 2% sucrose with ten-fold serial dilutions. White colonies were screened for double crossover strains by PCR verification. A correct knock-out strain was further verified by Southern blot and used in subsequent experiments.

For complementation of the knock-out strain, the upstream 500bp of the coding region for the AltRP operon (presumably incorporating the promoter region) along with the whole AltRP operon was cloned with C-terminal 3xFlag tag to AltRpsR and ligated to pML1342 as a complementation plasmid. The plasmid was transformed to ΔAltRP competent cells and plated on LB plates with 50μg/ml hygromycin. Recombinants were further verified by western blot in Sauton medium supplemented with different concentrations of zinc to check the expression from the operon.

### Generating endogenous AltRpsR/CanRpsR with different affinity tags strains

Upstream 500bp (US500) and downstream 500bp (DS500) of AltRpsR/CanRpsR stop codons were cloned. Different affinity tag sequences (3xFlag/3xHA) fused with different antibiotic expression cassette (Hygromycin/Zeocin) were generated by overlap PCR (3xFlag-Zeo oligo cassette was kindly provided by JHZ). The upstream, tag-antibiotic, downstream cassette were fused by Gibson assembly and cloned into pUC19 vector. pUC19 plasmid containing four different recombineering oligo cassette (AltRpsR-3Flag-Zeo, AltRpsR-3HA-Hyg, CanRpsR-3HA-Hyg and CanRpsR-3Flag-Zeo) were generated. *M. smegmatis* containing the plasmid pNitET-SacB-kan (58) was grown to OD_600nm_≈0.2 and 10μM isovaleronitril (IVN) was added to induce the expression of RecET, after inducing for 6h, competent cells were prepared by 3 washes in 10% glycerol. Oligos for recombineering were amplified from pUC19 series plasmid and 2μg DNA were transformed to the competent cells. Bacteria were plated on LB plates supplemented with either 50μg/ml hygromycin or 20μg/ml zeocin. Desired colonies were genotyped by PCR and further verified by Western Blot to test the expression of AltRpsR/CanRpsR by blotting against with either Flag or HA antibody.

### Western Blot to test zinc-switch phenotype

Strain A and Strain B were cultured in 7H9 medium to stationary phase (OD_600nm_≈3) and sub-cultured 1:500 to Sauton medium (zinc-free) supplemented with different concentrations of zinc sulfate. When bacteria reached stationary phase, bacteria were washed and re-suspended in TE buffer. The bacterial suspension was disrupted by beads-beating and lysate centrifuged at 17,950x*g*, 4°C for 5 min. Protein concentration was quantified by Bradford assay (Bio-rad) and same amount of protein were loaded onto 5%/12% SDS-PAGE gel. Proteins were further transferred onto PVDF membrane (Bio-rad). The membrane was blocked in 4% skim milk at room temperature (RT) for 1 h. Primary antibody was diluted 1:2000 and the blot was incubated at 4°C overnight. The membrane was then washed with TBS-T at RT and secondary antibody, diluted, 1:5000 rate and incubated at RT for 1 h. After washing with TBS-T, the image was developed by ECL Western Blotting Substrate (Pierce).

### Subunit profiling of *M. smegmatis*

For liquid culture, *M. smegmatis* strain was grown on Middlebrook 7H9 medium till OD_600nm_≈1 and collected by centrifuge at 4500x*g*, 4°C for 20 min. For plates, wild type *M.smegmatis* strain and ΔARP strain were plated on 7H10 plate or 7H10 plate supplied with 10μM TPEN and bacteria were collected by scraping. The bacterial pellet was washed and re-suspended with polysome buffer (50mM Tris-actate pH=7.2, 12mM MgCl_2_, 50mM NH_4_Cl). Bacteria were pulverized by beads beating or French Press, and the lysate were centrifuged at 18500x*g*, 4°C for 45 min. Cleared lysate were further loaded to a linear sucrose gradient (10%-50%) at Beckman SW41 rotor and centrifuged at 39,000 rpm, 4°C for 5 h. Gradients were fractioned with continuous A2_60_ measurement and fractions was collected for downstream analysis.

### Selective ribosome profiling with affinity- purified tagged AltRibosomes

Bacterial cultures of dual-tagged Strain A or Strain B were grown till OD_600nm_≈1 and treated with 100μg/ml chloramphenicol for 3 mins to arrest elongating ribosome before harvesting bacteria. Bacterial pellets were collect by centrifuging at 4500x*g*, 4°C for 10 min. The pellet was washed and re-suspended with ice-cold polysome buffer with minor recipe change. (50mM Tris-actate pH= 7.2, 12mM MgCl2, 50mM NH_4_Cl, 10mM CaCl_2_ and 100μg/ml chloramphenicol) The suspended bacterial suspension was dropped into liquid nitrogen and smashed into powder. The powder was smashed several times with repeatedly chilling with liquid nitrogen and was further pulverized with beads beating 4 to 5 times with chilling in liquid nitrogen between each step. After centrifugation at 18500x*g* for 45 min, the cleared lysate was transferred to a new tube supplied with 70U/ml MNase to digest ribosome-unprotected mRNA. The pre-digested lysate was laid over a 1M sucrose cushion and centrifuged at 30,000 rpm, 4°C for 20 h in Beckman 70Ti rotor. After ultracentrifugation, the ribosome pellet was washed and dissolved in polysome buffer. The dissolved ribosome solution was further digested with MNase at 4°C for 1 h. The digested ribosome fraction was incubated with either Flag resin or HA resin at 4°C overnight. After washing of the bound resin, the resin was eluted with 3xFlag/3xHA peptide to isolate the AltRibo (eluate), and the remaining flow-through was confirmed as consisting almost entirely of CanRibos.

The procedure for building deep sequencing libraries was followed as previously described (38) with minor changes. The detailed protocol is described as below. Ribosome-protected mRNA fragments were extracted by miRNeasy kit according to the manufacturer’s instructions and size-selected for 26-34nt fragment by marker oligo through 15% TBE-UREA PAGE. The corresponding region was sliced and smashed into pieces and soaked in 400μl RNA gel extraction buffer (300mM sodium acetate pH 5.5, 1mM EDTA and 0.25% SDS) and rotated at room temperature overnight. RNA was precipitated with 1.5μl GlycoBlue and 500μl isopropanol. The RNA was dephosphorylated with 1μl T4 PNK at 37°C for 1 h. RNA was extracted by isopropanol precipitation. Samples were then ligated with 0.5μl Universal miRNA cloning linker and 1.0μl T4 RNA ligase (truncated) at room temperature overnight. The ligation product was purified by isopropanol precipitation and visulized by 15% TBE-UREA PAGE. After slicing the corresponding ligated mRNA fragment, the samples were then subjected to reverse transcription with supersciptase III and RT-Primer. After incubating the reaction at 48°C, 30 min, the RNA template was further hydrolyzed with 2μl 1M NaOH at 98°C for 20 min. The RT product was precipitated with isopropanol and size-selcted with 15% TBE-UREA PAGE. The corresponding region was sliced and soaked in 400μl DNA gel extraction (300mM NaCl, 10mM Tris pH=8, and 1mM EDTA) and rotated at RT overnight. The cDNA was purified by isopropanol precipitation and circularized with 1μl CircLigase and incubated at 60°C for 1 h. The circularized cDNA template was incubated with 1μl 10mM Biotinylated rRNA subtraction oligo pool mix. The Oligo sequence was adopted from another study (39) with additional oligos newly designed according to our preliminary ribosome profiling data (Table S2). The mixture was denatured for 90s at 100°C and then annealed at 0.1°C /sec to 37°C. This was then incubated at 37°C for 15 min. 30μl streptavidin C1 DynaBeads was added to the mixture and incubated for 15 min at 37°C with mixing at 1000 rpm. Then the mixture was placed on a magnetic rack to isolate the beads, the supernatant was transferred to a new tube and cDNA purifed by isopropanol precipitation. The subsequent cDNA template was used for library amplification with Phusion high-fidelity DNA polyermase for 12-14 cycles. PCR products were purifed from 8% TBE PAGE gel. The corresponding bands were sliced and soaked in DNA gel extraction buffer and rotated at room temperature overnight. The PCR products were purified by isopropanol precipitation and the concentration and quality were measured with Agilent 2100 Bioanalyzer. Libraires were sequenced on the illumina X-ten sequencer.

### Large-scale tagged ribosome purification for biochemical analysis

Bacterial cultures (~30L) were grown to around OD_600nm_≈1 and bacterial pellets were collected by centrifugation at 4500x*g*, 20 min, 4°C. Pellet was washed and re-suspended in polysome buffer. Bacteria were disrupted by French Press and centrifuged at 18500x*g*, 4°C for 1h to collect supernatant. MNase was added to the cleared lysate at a concentration of 70U/ml and digested at 4°C overnight. The digested lysate was loaded onto 1M sucrose cushions and centrifuged at 30,000 rpm, 4°C for 20 h. After centrifugation, the supernatant was discarded, and the ribosome pellet washed with polysome buffer and dissolved in polysome buffer with gentle shaking at 4°C. The crude ribosome suspension was further digested with MNase at 4°C overnight. After secondary digestion, the ribosome preparation was dialyzed against HEPES-polymix buffer (59, 60). The ribosome preparation was incubated with anti-Flag M2 beads 4°C overnight. After binding with M2 beads, the flow-through was collected as CanRibo and AltRibo was collected by eluting with Flag peptide. Each fraction, comprising total Ribo, CanRibo (flow through) and AltRibo (Eluate), were probed by Western blotting for quality check and purity prior to further analysis.

### *In vitro* 70S initiation complex formation

Initiation complex was formed by incubating 70S ribosome variants as described above (0.2 μM), [^3^H]fMet-tRNA^fMet^ (0.5 μM), XR7mRNA (AUG-UAA) (0.5 μM), initiation factors (0.5 μM) and GTP (1 mM) at 37°C for 15 min in HEPES-polymix buffer (59). The reactions were thereafter filtered through BA-85 nitrocellulose membranes and washed with 10ml of ice-cold HEPES-polymix buffer (pH 7.5). The radioactivity retained on the filter was measured in a Beckman LC6500 scintillation counter.

### In vitro initiator tRNA binding

BODIPY labelled Met-tRNA^fMet^ (0.1 μM) was rapidly mixed in stopped flow with 70S ribosomes (1 μM) pre-incubated with XR7mRNA (AUG-UAA) (0.5 μM). BODIPY fluorescence was excited at 575 nm and the change in fluorescence signal was monitored in real time after passing through a 590 nm cut-off filter. The time course data was obtained by averaging 3-5 individual traces and fitted with single exponential model.

### In vitro di-peptide assay

For Met-Leu dipeptide formation, an initiation complex was formed with mycobacterial 70S ribosome variants (1-5 μM), AUG-CUG-UAA XR7 mRNA (20 μM) and [^3^H]fMet-tRNA^fMet^ (1 μM), initiation factors (2 μM) at 37°C. In parallel, an elongation mix was prepared which contained *M. smegmatis* EF-Tu (20 μM), EF-Ts (20 μM), tRNA^Leu^ (100 μM), Leu amino acid (0.5 mM), Leu-tRNA synthetase (1unit/μL) and GTP (1mM) at 37°C. Both initiation complex and elongation mix containing energy pump ATP (1mM), EP (10 mM), PK and MK (Sigma). The reaction was started by mixing equal volumes of the initiation complex and elongation complex at 37°C and was quenched after 10 s by adding 17% formic acid. The peptides were isolated from the ribosome complex by KOH treatment and further analyzed by HPLC equipped with a radioactivity detector.

### Read preparation and alignment

The *M. smegmatis* mc^2^-155 reference genome assembly (ASM1500v1) downloaded from Ensembl Bacteria Release 32 (https://bacteria.ensembl.org/) was used for all analyses. For RNAseq, we performed quality control and trimmed adaptors using the FastQC and FASTX-Toolkit. For ribosome profiling, after quality control and adaptor removal, reads less than 25nt long were discarded and longer than 36nt were trimmed to 36nt. Next, ribosome profile and RNAseq reads that aligned to rRNA and tRNA were removed using bowtie2 (61). We aligned the remaining reads to the genome using bowtie2 with “sensitive-local” option. The mapped reads were normalized to reads per kilobase million (RPKM) using total number of mapped reads.

### A-site assignment

For each read, the A-site’s position was inferred by taking the first nucleotide that was 12nt upstream of the 3’end of the read (62). Based on this, we generated a signal track of A-sites for all transcripts. ORFs that had been annotated in the reference were denoted as annotated ORFs (aORFs).

### Polarization of aligned Ribosome Profile reads

The polarity score was calculated on the basis of aligned A-site signal track by adapting a previous method (40). The observed polarized reads from AltRibos were not due to ambigous artifacts that might be caused by peaks around the start/stop codons, since the same result was observed whether or not the first and last 15nt of the coding region were included in the analysis. aORFs with more than 50 reads in coding sequences were plotted. In this study, we calculated the polarity score under AltRibo, CanRibo as well as total ribosome population. Here we defined the concept of polarity shift by taking total ribosomes as the background input and subtracting the polarity score of background from the score of AltRibo-enriched/CanRibo-enriched populations.

### Differential ribosome codon reading

The calculation of codon occupancy was adapted from a previous method (41). For each aORF, only in-frame ribosome profiling fragments (RPFs) were included to study the frequencies of all codons. For each coding gene, its codon density was calculated by normalizing the observed RPF frequencies by the total frequcies of codons. Later we included genes with at least 100 RPFs as input and obtained an averaged codon density for each codon. Here we denoted *T*_*c*odon_*i*__. as the averaged codon occupancy of *c*odon_*i*_ for AltRibo or CanRibo population while *R*_*c*odon_*i*__ as the averaged codon occupancy of *c*odon_*i*_ for reference population, in this case, the total ribosomes. To better clarify the difference between AltRibo and CanRibo population, we further defined codon occupancy shift as follows:

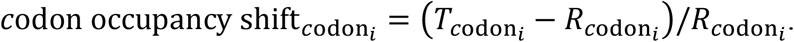

Significantly different codon usage was detected by comparing the codon density of all studied aORFs under test conditions (AltRibo/CanRibo) against the corresponding density under the reference condition (total ribosome input) (t-test *p value* <0.05 and shift > 0.05 or shift < −0.05). Additionaly, for each individual aORF we correlated its codon density from AltRibo/CanRibo population with that from total ribosome population (Pearson correlation was used in this study).

### Differential codon abundance

In this study, we used the concept of codon abundance to measure the occurance of specific codons in the 50 codons within the 5’ or 3’ of the coding region. The calculation of codon abundance followed a previous study (63). To better demonstrate the difference, the final results were presented in the following form:

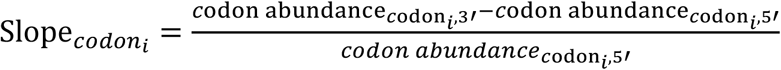

Slope*_codon_i__* represents the slope of *codon_i_* in which *c*odon abundance_*c*odon_*i*_,3′_ represents the codon abundance of *codon_i_* in the 3’ region while *c*odon aboundance_*c*odon_*i*_,5′_ represents the codon abundance of *codon_i_* in the 5’ region.

### Differential translation analysis

Since AltRibo, CanRibo and total ribosomes were collected from the same population of bacteria, to identify differential translated genes we used edgeR on the corresponding ribosome profiling datasets (64). During analysis, replicates were included. Differentially translated genes were identified by log_2_fold change (FC) > 0.5(up-regulated) or log*2*FC< −0.5(down-regulated) and P value < 0.05. In this study, the datasets of total ribosome were treated as the control and compared against AltRibo or CanRibo to identify differentially translated genes.

### Statistical Analysis

Appropriate statistical tests are detailed within each protocol and in the figure legends.

## Data availability

Ribosome profiling and RNA-seq data presented in this study is available with GEO accession number GSE127827.

## Acknowledgements

We thank Jianhuo Fang from the Tsinghua genomic and synthetic biology core for help in preparing the ribosome profile libraries. We would like to thank Eric Rubin and Hesper Rego for reading and commenting on the draft manuscript. This work was in part funded by grants from the Bill and Melinda Gates Foundation (OPP1109789) and from the National Natural Science Foundation of China (31570129) and start-up funds from Tsinghua University to BJ and from the Swedish Research Council (2018-05498, 2016-06264) and Cral-Tryggers Foundation (CTS 18:338) grants to SS. BJ is an Investigator of the Wellcome Trust (207487/B/17/Z).

## Author Contributions

YXC and BJ designed the experiments and interpreted the data. YXC performed the majority of the experiments, with XG performing the biochemical experiments. ZYX and YXC analyzed the ribosome profile data. ZJL, SS and BJ supervised research. YXC and BJ wrote the manuscript with input from the other authors.

