## Supplementary material for "Selective translation by alternative bacterial ribosomes": Supp Figures

### Figure S1

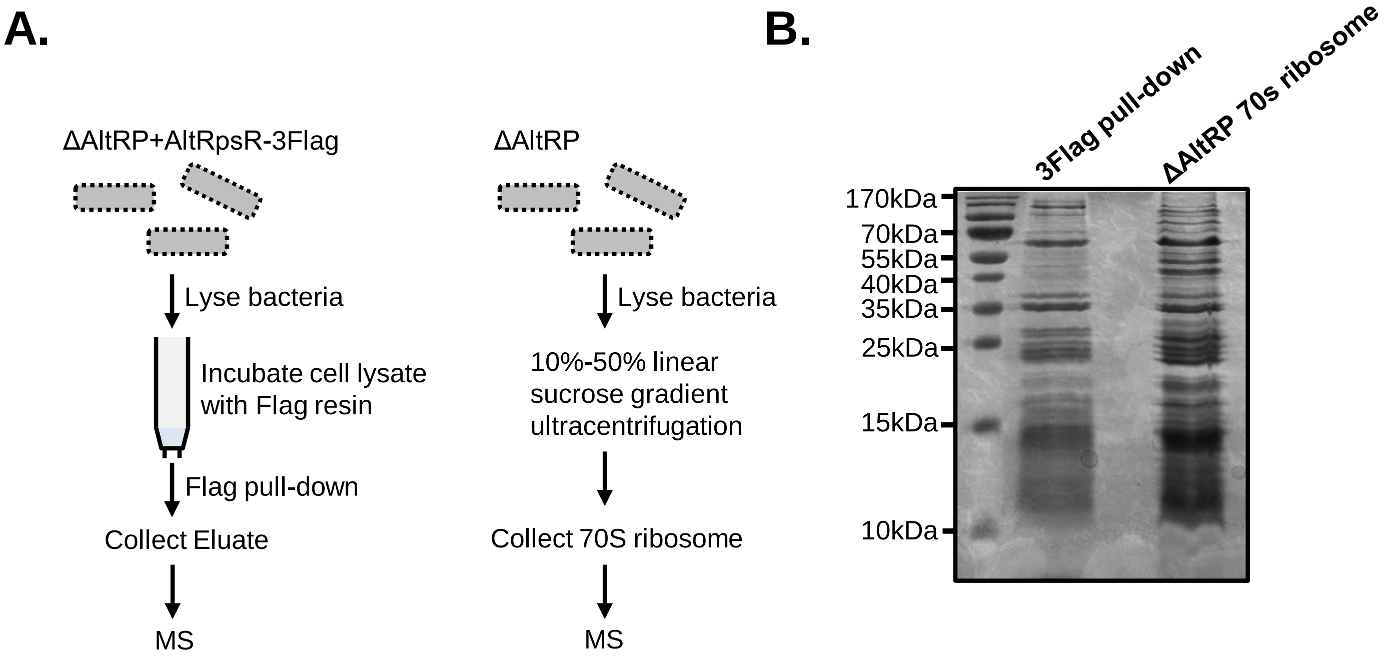


**Figure S1**. **Affinity purification of tagged AltRpsR co-immunoprecipitates the ribosome.**
(A) Cartoon representing the work-flow for immunoprecipitation of FLAG-tagged AltRpsR (left panel) and total ribosomes (right panel). (B) Coomassie-stained SDS-PAGE gels of anti-FLAG eluate and 70S ribosomes from sucrose-density centrifugation as shown in (A).

### Figure S2

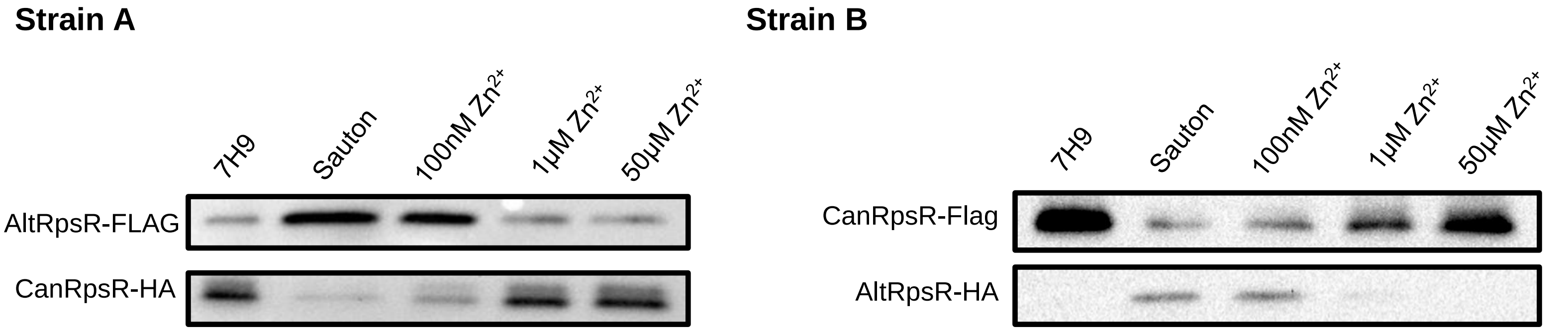


**Figure S2. AltRpsR expression is negatively regulated by zinc.**
Anti-FLAG and anti-HA Western blots of cell lysate (top panel) and of ribosome pellets (bottom panel) in complete 7H9 medium (7H9) and in standard Sauton's medium (Sauton) or Sauton's medium supplemented with zinc sulfate (at indicated concentrations) of Strain A (a) and Strain B (b).

### Figure S3

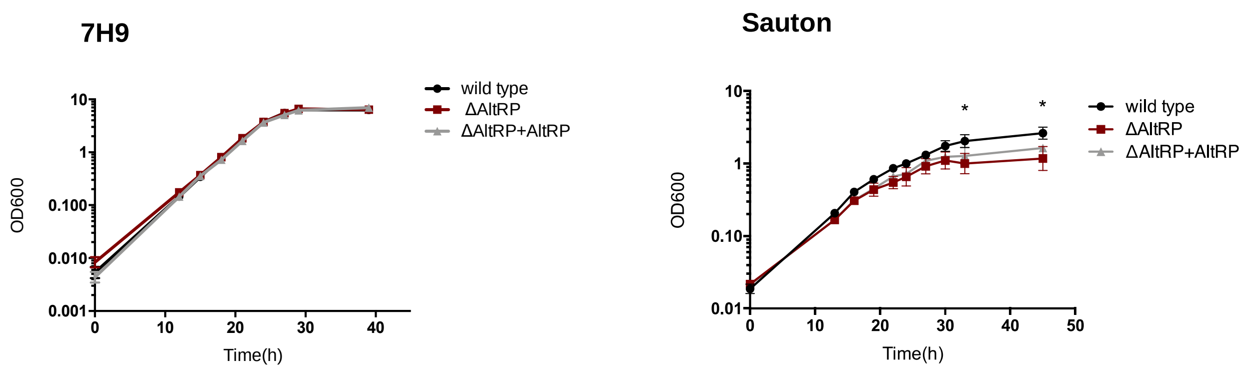


**Figure S3. Strain *M. smegmatis*-ΔAltRP has a growth defect in zinc-poor medium.**
Growth curves (as measured by $OD_{600nm}$) for wild-type *M. smegmatis*, *M. smegmatis*-ΔAltRP (ΔAltRP) and *M. smegmatis*-ΔAltRP::AltRP (ΔAltRP+AltRP) in zinc-replete complete 7H9 medium (left panel) and relatively zinc-deplete Sauton's medium (right panel). Three biological replicates +/- SD are plotted

### Figure S4

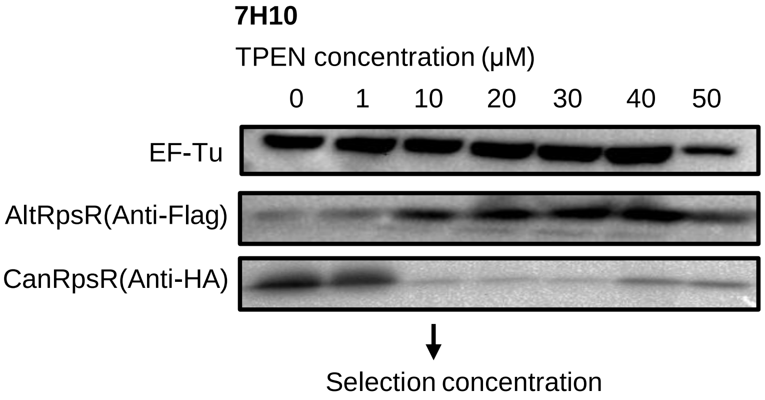


**Figure S4. 10 µM TPEN is sufficient to alter the expression profile of AltRpsR and CanRpsR.**
Western blot against CanRpsR (anti-HA) and AltRpsR (anti-FLAG) for wild-type *M. smegmatis* scraped from 7H10-agar plates supplemented with increasing concentrations of TPEN. Antibody against EF-Tu was used as a loading control.

### Figure S5

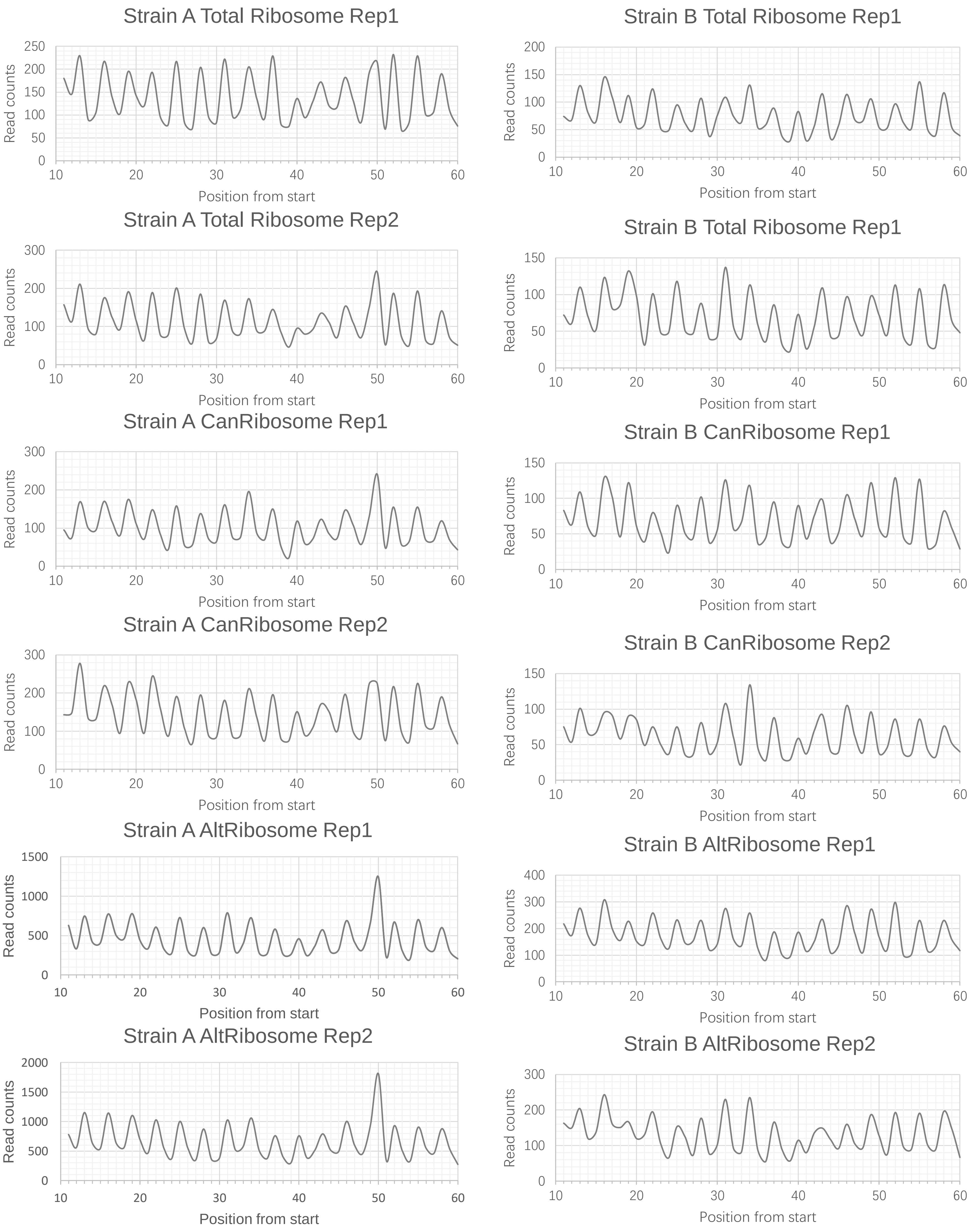


**Figure S5. Ribosome profile reads show clear 3 nucleotide periodicity.**
Representative ribosome profile 28nt reads from each replicate from Strains A (A) and B (B) used in this analysis.

### Figure S6

**A.**

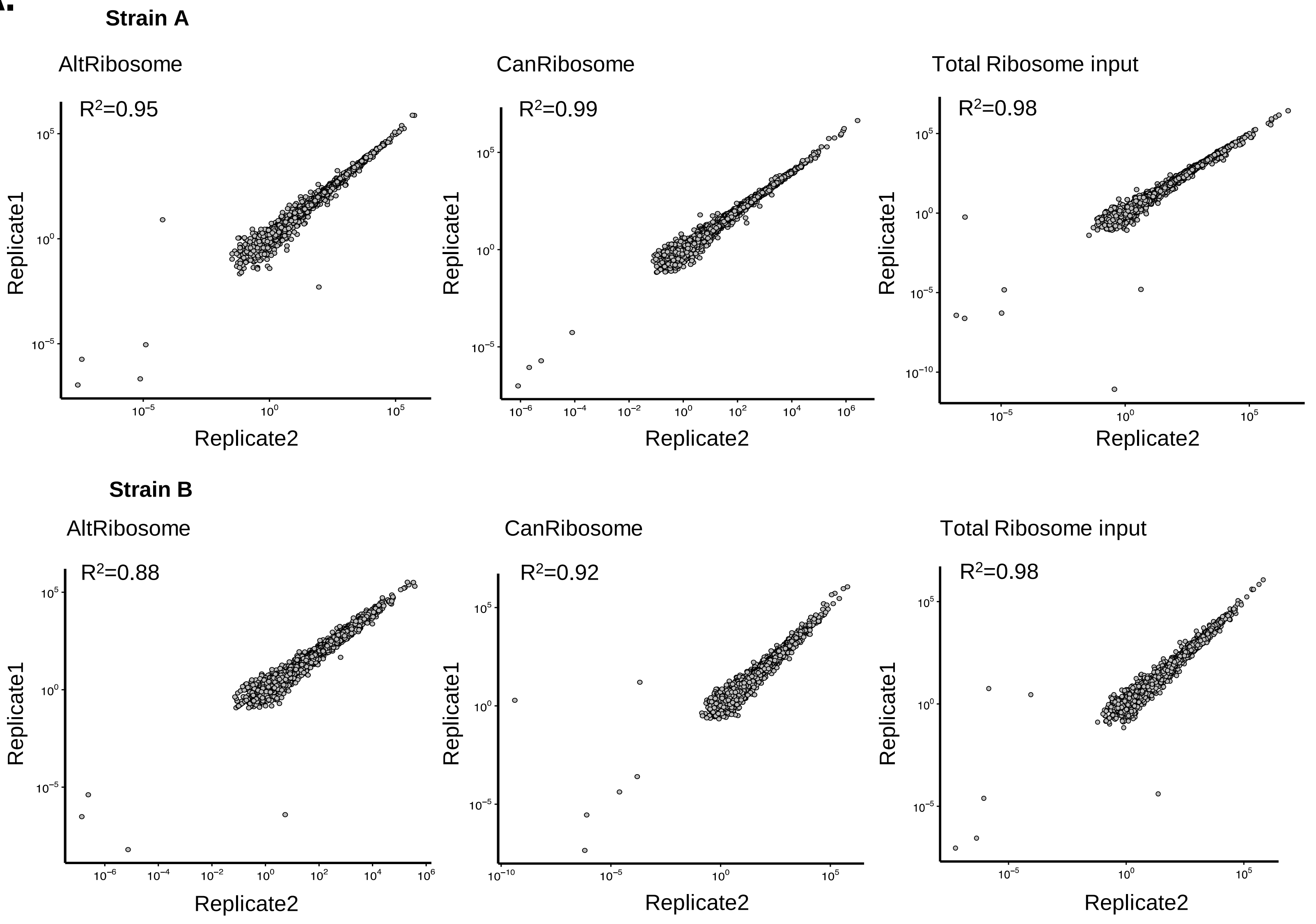


**B.**

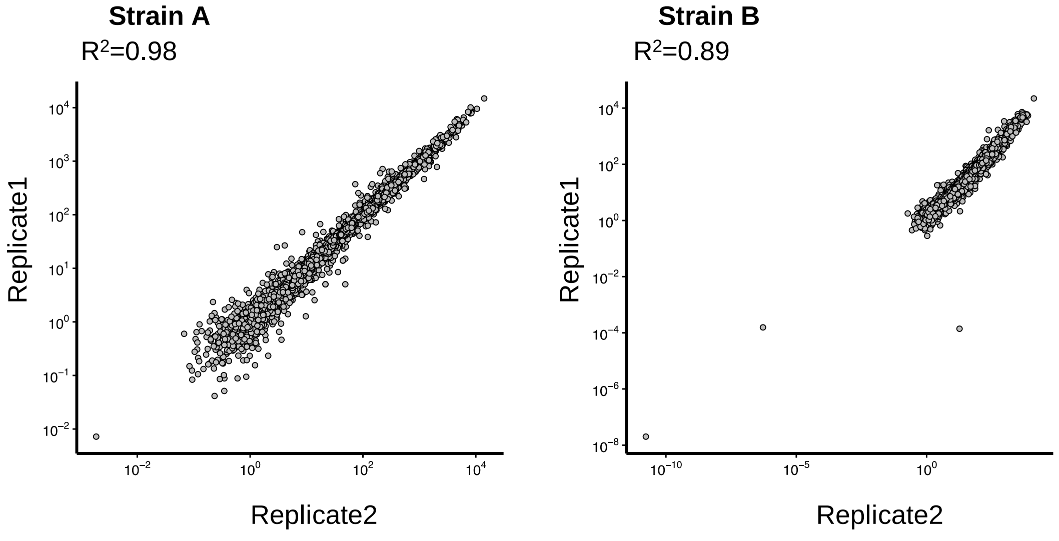


**Figure S5. Biological replicates are highly correlated.**
(A) Pearson correlation analysis of selective AltRibo, CanRibo and total ribosome profiles of the two biological replicates from each strain used in this study. (B) Pearson correlation analysis of paired RNAseq of the two biological replicates from each strain used in this study.

### Figure S7

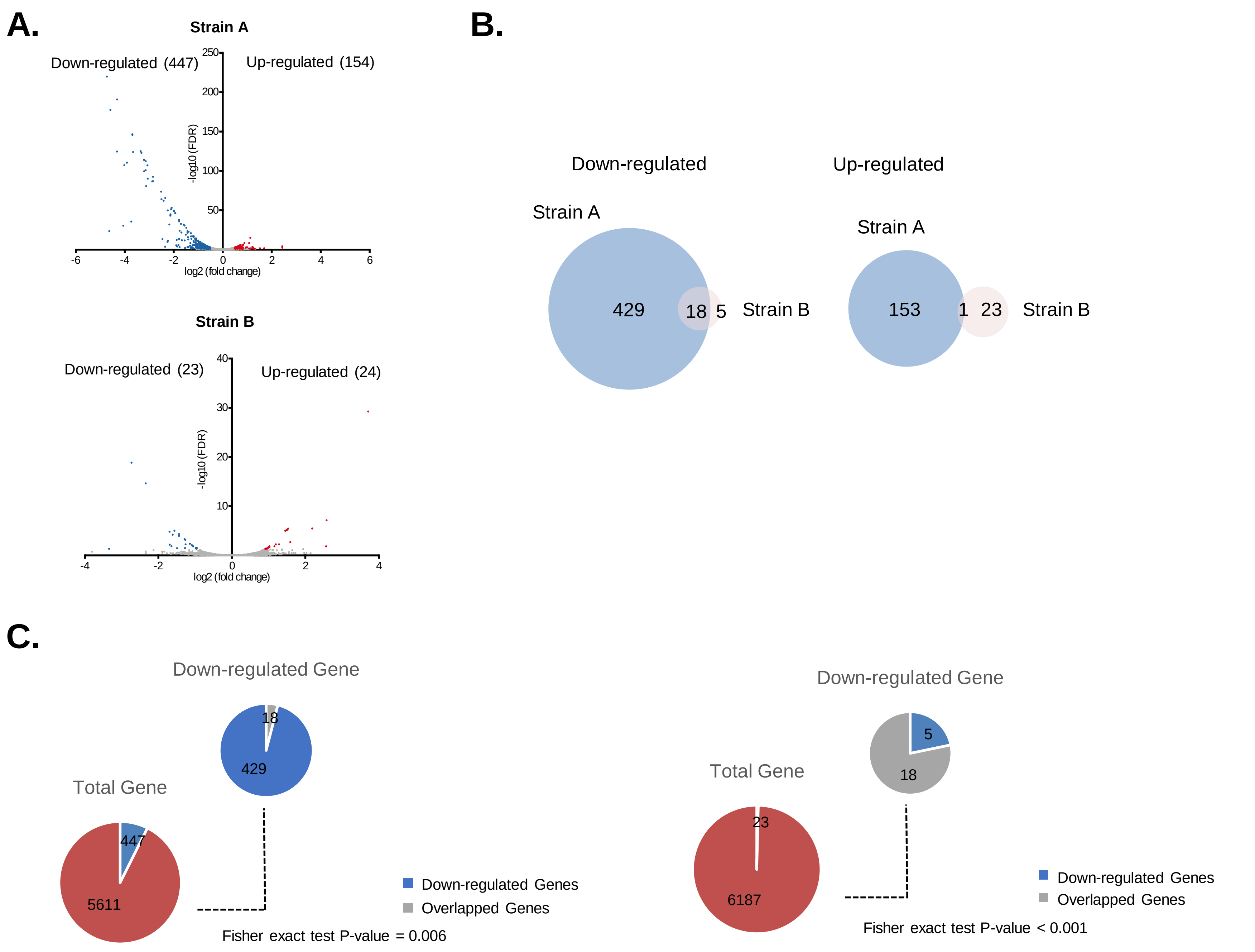


**Figure S7. AltRibo down-regulated genes have a high degree of overlapping in Strain A and Strain B.**
(A) Normalized ribosome profile read density for AltRibos compared with total ribosome input in Strain A and Strain B. Significantly upregulated ($Log_2$ fold-change (FC) > 0.5, False discovery rate < 0.05) and down-regulated genes ($Log_2$ fold-change (FC) < -0.5, False discovery rate < 0.05) are represented by red and blue dots respectively. (B) Venn diagram show the overlapped genes in both down-regulated and up-regulated gene categories in both Strain A and Strain B. (C) The 18 overlapped down-regulated genes are significantly enriched in both Strain A and Strain B, as measured by Fisher's exact test.

### Figure S8

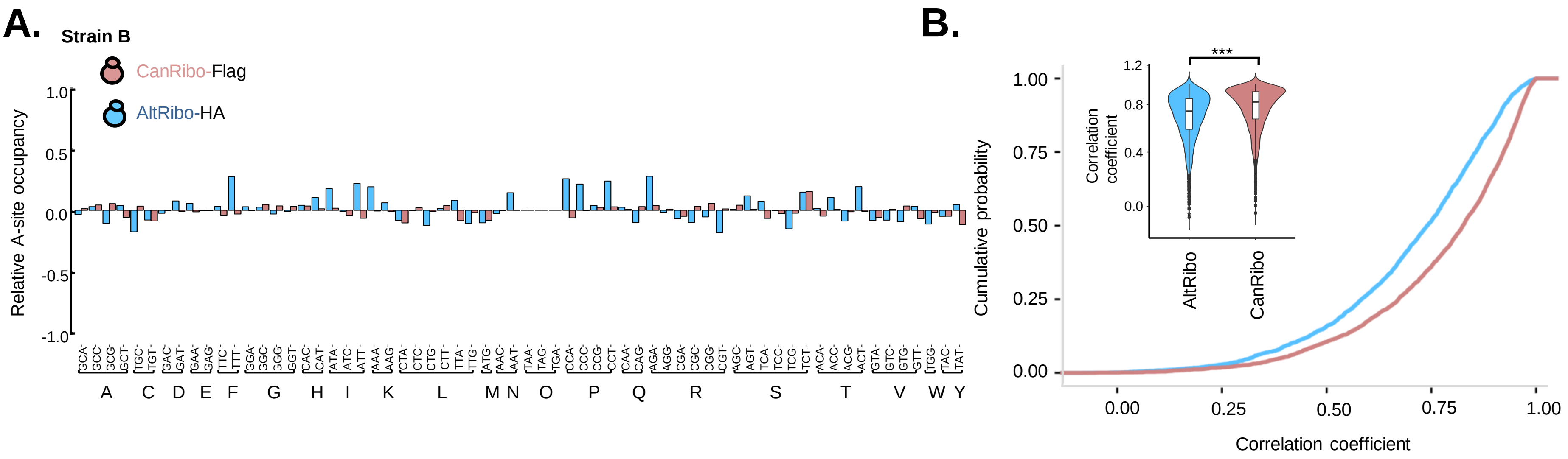


C.

| AltRibosome high occupancy codons | AltRibosome low occupancy codons |
|---|---|
| AAA(K) | ACG(T) |
| AAG(K) | CAG(Q) |
| AAT(N) | CGC(R) |
| ACC(T) | CGG(R) |
| ACT(T) | CGT(R) |
| AGA(R) | CTG(L) |
| ATT(I) | GCG(A) |
| CCA(P) | GTC(V) |
| CCC(P) | GTG(V) |
| CCT(P) | TCG(S) |
| GAA(E) | TGC(C) |
| TTT(F) | TGG(W) |

**Figure S8. Alternative ribosomes have distinct codon preferences compared with canonical ribosomes.**
Analysis as per Fig. 3c-d, but for Strain B. (A) The relative A-site codon occupancy for all 61 sense codons for AltRibos and CanRibos in Strain B compared with total ribosome input. (B) Cumulative probability and violin (inset) plots for correlation between codon occupancy for AltRibos and CanRibos compared with total ribosome input from Strain B. *** $p<0.001$ by Student's t-test. (C) Combining data from Strain A and B, AltRibos have 12 high and 12 low occupancy codons compared with CanRibos.
